# NMR spectroscopy of a single mammalian embryo

**DOI:** 10.1101/2021.07.26.453772

**Authors:** Giulia Sivelli, Gaurasundar M. Conley, Carolina Herrera, Kathryn Marable, Kyle J. Rodriguez, Heinrich Bollwein, Mateus J. Sudano, Jürgen Brugger, Andre J. Simpson, Giovanni Boero, Marco Grisi

## Abstract

The resolving power, chemical sensitivity and non-invasive nature of NMR has made it an established technique for *in vivo* studies of large organisms both for research and clinical applications. These features would clearly be beneficial at the nanoliter scale (nL), typical of early development of mammalian embryos, microtissues and organoids, the scale where the building blocks of complex organisms could be observed. However, the handling of such small samples (about 100 micrometers) and sensitivity issues have prevented the widespread adoption of NMR. Recently we have shown how these limitations can be overcome with ultra-compact single-chip probes. In this article we show that such probes have sufficient sensitivity to detect and resolve NMR signals from individual bovine pre-implantation embryos. In less than 1 hour these spherical samples of just 130-190 micrometers produce distinct spectral peaks, largely originating from lipids contained inside them. We further observe how the spectral features, namely the peak intensities, vary from one sample to another despite their optical and morphological similarities.

## Introduction

The use of Nuclear Magnetic Resonance (NMR) for imaging and spectroscopy is nowadays routine in clinics, hospitals, and R&D centers. NMR is recognized to be a key enabler of many life-saving applications and the gold standard for non-invasive *in vivo* analysis^1-4^. In terms of imaging, besides unique abilities for physiological contrast, what separates NMR from other methods (e.g. X-ray imaging) is the non-ionizing radiation required, which makes it possible to observe intact lifeforms even during long analysis procedures. The static magnetic field alone has no, or extremely small, effects on cell growth and genetic toxicity regardless of the magnetic field density^5^. Furthermore, given a gyromagnetic ratio of 42.58 MHz/T for ^1^H nuclei, at a field strength of about 7 T (210’000 times stronger than the Earth’s magnetic field) the resonance takes place at a frequency of just 300 MHz. Irradiation with wavelengths of such low energy (i.e. radio waves) is too weak to break or even vibrate molecular bonds, therefore avoiding genetic changes and preserving the chemical environment in its true *in vivo* and unaltered state (key for metabolic functioning). The main concern related to NMR safety is localized radio frequency heating, which must be maintained in the physiological range of the organism being analyzed. This is the case when the Specific Absorption Rate (SAR) remains below a few W/kg during the clinical examination^6,7^. Recently, full MRI operation at 7 T was approved for human use by the Food and Drug Administration (FDA), making it the highest field that can support clinical application to date (Magnetom Terra, Siemens Healthineers)^8^.

However, it is the spectroscopic capabilities (rather than imaging) of NMR that truly set it aside as a unique analysis approach, and arguably, hold the most potential for diagnosis and scientific discovery. NMR, without use of libraries, can solve the structure of compounds (both crystalline and non-crystalline) *de-novo*. It is the only molecular-level tool that can provide non-invasive and comprehensive information as to the structure and interactions inside living organisms. For example, the first metabolome of a living organism (Daphnia Magna, essential keystone species used in toxicity testing) was recently assigned in 2020^9^. Further, NMR holds unique capabilities to study non-covalent binding^10^, including mechanisms and receptors, essential life processes and pharmaceutical development, toxicity and fate.

Due to the lack of non-invasive biochemical assays for exceedingly small samples such as mammalian embryos, NMR represents a valuable candidate to bring a substantial advantage at the microscopic scale, and would provide the basis for R&D studies and further clinical applications that are currently out of reach. For example, stem cells are now known to be the foundation of development in plants, animals and humans and understanding the chemistry behind these processes offers a transformative approach for future medicine. However, the primary and fundamental obstacle to micro-NMR is the unfavorable scaling of signal as the volume is reduced. To overcome this issue, a micro-coil is needed that approximately matches the sample size^11^. This justified many efforts for the development of NMR micro-coils^12,13^, delivering spectroscopy studies on biological samples such as eggs and embryos of insects^14,15^, amphibians^16,17^, fish^18-20^, whole microorganisms^21,22^, and giant neurons of sea snails^23,24^ down to a scale of approximately 10 nanoliters (Fig. 1).

**Figure 1:**
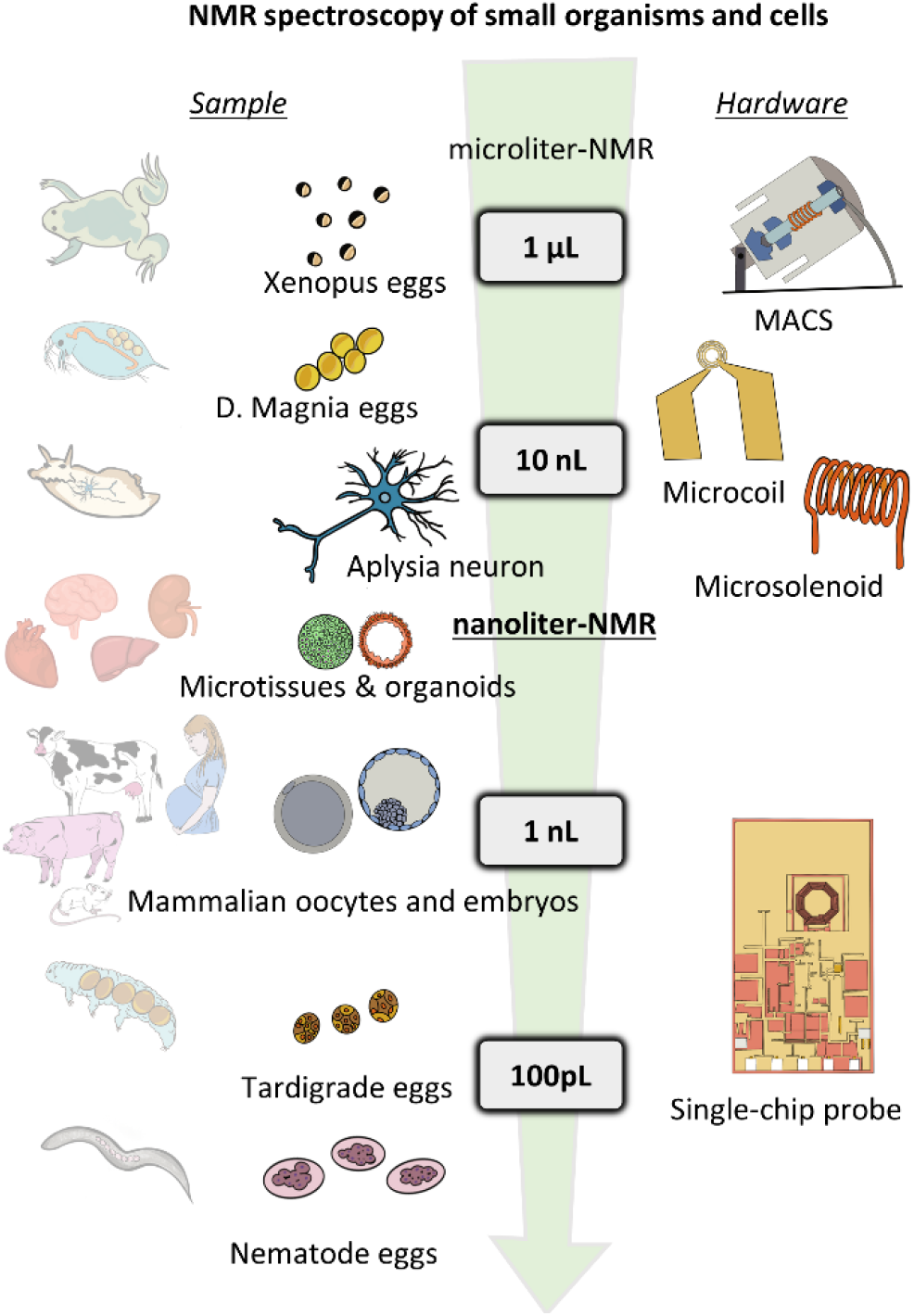
NMR spectroscopy of small organisms and cells. Microliter, nanoliter, and subnanoliter single biological entities (eggs, embryos, oocytes, and neurons from different animals) investigated by NMR spectroscopy using dedicated inductive detectors (magic-angle spinning solenoids, planar microcoils, micro-solenoidal coils, and single-chip probes with micro-coil and electronics co-integrated).

These studies have shown the potential of microscopic NMR, but smaller samples remained out of reach mainly due to sample handling issues and fundamental geometrical constraints (the connections from the micro-coil to the electronics remain bulky and introduce parasitic sensing regions)^25^. Incidentally, sample volumes of about 1 nanoliter would open huge potential for novel applications: at such scale we indeed find organoids, 3D cell cultures, and early stage animal and mammalian embryos (Fig 1) for which a non-invasive biochemical assay is still lacking. These same specimens are at the core of several drug development and toxicity studies, as well as personalized medicine. Other non-invasive methods based on alternative spectroscopic techniques, such as Fluorescence Lifetime Imaging, are being developed for early stage embryos, but the analytical range remains limited to the few molecules that exhibit autofluorescence^26-28^.

More recently, our group has shown that single-chip micro-probes in combination with 3D micro-printing can overcome critical issues and expand the reach of *in vivo* NMR to the nanoliter scale. With this approach we were able to show that is possible to perform spectroscopy of microorganism eggs^29,30^ and human microtissues^31^ in the nanoliter range and below. Coincidentally, all mammalian species start to develop at the scale of about 100 microns: such is the size of their oocytes until developing into blastocysts a few days after fertilization. In this short communication we show the first data obtained from NMR spectroscopy of single pre-implantation stage cattle embryos, the bovine model being one that closely reflects human embryo development and reproductive physiology^32^. In these experiments we use bovine embryos obtained through *In Vitro* Fertilization (IVF) techniques, produced in UZH (Zurich). The embryos are then transported cryopreserved to our laboratory at EPFL, where they are measured shortly after the thawing process (see Material and Methods).

We should note that conventional fluid NMR spectroscopy methodologies have been used previously to investigate the implantation potential of individual human embryos^33,34^. In these studies it is the liquid medium where the embryo is cultured that is analyzed. Such liquid NMR measurements naturally provide higher spectral resolutions and reduce concerns over sample handling. However, the information that can be obtained is inherently more limited. One can only infer biochemical information from the embryo indirectly, through measurements of uptake and expulsion of compounds from and into the medium. Additionally, the embryo itself has a volume (∼ nL) that is about 10000 times smaller than that of the culture medium (∼10 µL), making accurate quantification very challenging^34^. In contrast, the micro NMR method here proposed targets the biological sample directly, providing expanded access to endogenous compounds.

## Materials and methods

Experiments and protocols were approved by the Life Science biosecurity unit of the École Polytechnique Fédérale de Lausanne (EPFL) and carried out in accordance with the experimentation guidelines of the institution.

### Microscopic NMR set up and experimental details

Fig. 2A shows a schematic of the setup consisting of a Bruker 300 MHz magnet, a nano-NMR probe designed with a plug & play interface for the microchip-based NMR sensors, and an adapted console for signal generation and acquisition. The details of the microchip-based NMR probe are reported in Ref [31]. Fig. 2B shows a photograph of the microchip-based sensor, consisting of a microsystem enclosed in a 3D printed plastic container holding the cell culture medium used during these experiments. The container can be opened for easy access to the sample chamber via a micro-pipette, and it is designed to seal the chamber during the operation in order to reduce evaporation. The sample is easily placed in the sensing region via a micro-pipette with the aid of a stereo-microscope in a dust free environment. The loaded sensor is then plugged into the receptacle, inserted in the magnetic field, and the experiments run for 50 minutes.

**Figure 2:**
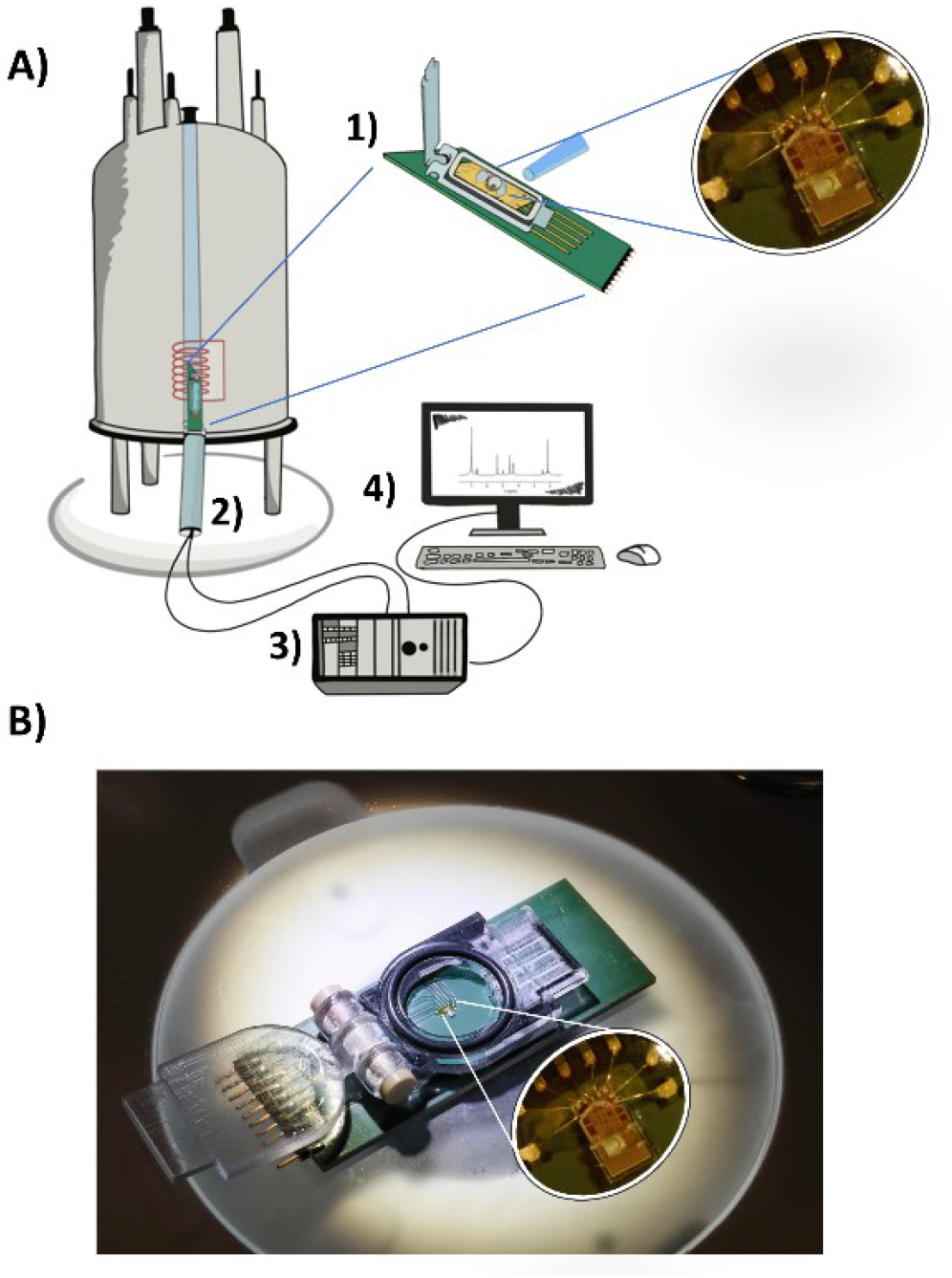
nanoliter-NMR setup and microchip based sensor. **A)** NMR setup: 1-micro NMR sensor 2-custom made sensor loader, 3-console managing triggers, RF, and data acquisition, 4-GUI and software processing. **B)** Photograph of the microchip-based sensor (width: 2.5 cm, length: 6 cm), consisting of a microsystem enclosed in a 3D printed plastic container holding the cell culture medium used during these experiments (protocol in material and methods). Similarly to Ref. [31], where more details are described, this system mounts a micro-structure to hold the sample during the experiments. In this case the dimensions have been adapted to hold spherical samples of approximately 160 μm.

### NMR experimental details

NMR experiments were performed in the 54 mm room temperature bore of the Bruker 7.05 T (300 MHz) superconducting magnet. The electronic setup is similar to the one described in detail in Ref. [35]. In this realization, the RF sources, DAQ, and TTL pulses are generated by a modified Tecmag Scout console, whose GUI is used to manage the experimental parameters. Two different frequencies are used in transmit and receive mode with a resulting down-conversion at about 5 kHz. Transmit frequency is fixed at the Larmor frequency. All experiments were performed with a repetition time of 400 ms, a pulse length of 20 μs, and an acquisition time of 140 ms with 2048 data points.

The final result is based on the averaging of 7500 scans, for a total experimental time of 50 minutes. Prior measurements the magnetic field is shimmed with a probe without sample (i.e., a sample of culture medium) and left untouched for the day. The data processing is automated with a custom Matlab software, all spectra are processed with the same algorithm as follows: (1) the time domain signal is post-processed by applying an exponential filter with decay time of 50 ms; (2) the spectrum is computed with an FFT; (3) Spectra are phased to minimize the negative points of the real part from 1 to 7 ppm; (4) two linear fits are used to remove the baseline in the spectral region using baseline data from −5 to 0 ppm and from 9 to 12 ppm; (5) Steps 3 and 4 is iterated; (6) The frequency axis is transformed to ppm units, and the maximum value of the water peak is defined at 4.75 ppm.

### Sample delivery

Ovaries retrieval and processing, oocytes isolation, IVF, Embryo Culture and cryopreservation were carried out at UZH (Zurich). Frozen embryos loaded in cryopreservation straws (5 embryos per straw) were delivered in a liquid nitrogen dry shipper to the laboratory and immediately transferred to a liquid nitrogen biobank.

### Embryo thawing

Embryo thawing was carried out 3 hours prior to the NMR measurement following protocols previously optimized at UZH. On the day before thawing, *in vitro* culture (IVC) media dish were prepared by filling 4 wells of a 4-well IVF dish with 500 μl BO-IVC medium (IVF Bioscience) per well. The IVC plate was incubated overnight in a humidified incubator with 5% CO_2_, 6% O_2_, and balance of N_2_ at 38°C in order to allow media to equilibrate. On the day of the experiment the thawing dish was prepared as follow: 2 wells of a 4-well IVF plate were filled with 500 μl each of thawing solution (Hepes-buffered TCM-199 with 10% fetal bovine serum and 0.5 M Sucrose) and the remaining 2 wells were filled with 500 μl each of holding solution (Hepes-buffered TCM-199 with 10% fetal bovine serum). The thawing dish was maintained on a thermal plate set at 38 °C. The straw containing the embryos to be thawed was placed in a styrofoam box filled with LN_2_. Then the straw was air thawed for 3 s and transferred into a water bath set at 24 °C for 30 s. The straw was removed and wiped with a sterile gauze. The plastic plug was removed from one end of the straw and the open end held over a small petri dish containing 500 μl of warming solution while cutting the opposite end to let the content of the straw freely flow into the warming solution. The embryos were quickly transferred to the first well of warming solution and left for 5 minutes and the process was repeated for the subsequent wells of warming and holding solution. The embryos were then transferred in the first well of pre-equilibrated IVC medium (BO-IVC, IVF Bioscience, UK) and washed 5 times by gentle pipetting. Embryos were transferred in the second well of pre-equilibrated IVC medium and placed into the incubator with 5% CO_2_, 6% O_2_ and balance of N_2_ at 38 °C for 3 hours before proceeding with the NMR measurement.

### Sample handling

NMR chamber was loaded with 2 mL of pre equilibrated BO-IVC medium and checked for possible air bubble presence under a stereomicroscope. When present, air bubbles were removed by gently pipetting. A single embryo was removed from the culture dish and placed in the sensor using a Cook Medical 300 μm tip and pipette. The NMR box was then closed and the sensor connected to a manual slide & lock mechanism to position the device into the NMR magnet (see Ref. [31]). NMR measurements were performed for a total time of 50 minutes per embryo. A picture of each embryo before and after NMR measurement was taken using a Drawell stereomicroscope to further confirm correct embryo positioning inside the microwell on the chip.

## Results

In order to obtain a first proof of concept for pre-implantation embryo selection we decided to run this study on *in vitro* produced pre-implantation bovine embryos arrested at different developmental stages. This choice arose from the necessity to be able to analyze them at different and fixed developmental stages in order to investigate microchip NMR technology’s discrimination potential. In addition, we are conscious that the current version of the device is not yet able to keep a stable internal environment suitable for live embryos analysis therefore, we could have run into the possibility of not knowing if the embryo was previously arrested or if the exposure to the device was the arresting cause. Despite these limitations, we were still able to establish a proof of concept. An upgrade version of the technology is being developed to analyze live samples.

The bovine is one of the mammalian models that better reflects human embryo development and reproductive physiology^32^. Bovine, alongside porcine embryos, are known to contain an increased lipid content when compared with mouse and human embryos, which suggests a direct relationship between lipids and the interval between ovulation and implantation in mammals^36^. It is well documented that lipids participate in key biological process and cellular pathways, although a complete understanding of the lipid importance for early development and embryonic competence is still lacking. Here we selected samples ranging from 2-cell stage up to morulae, mainly because of their spherical shape and relatively constant size across this development range, making it possible to carry out all measurements with a single sensor design.

Fig. 3B shows the microsystem loaded with a bovine embryo, where the sample is held in place by a micro-structure as previously shown in Ref. [31]. The micro-structure holding the embryo is designed so that the sample is held in the most sensitive region of the device, entirely determined by the on-chip micro-coil. Fig. 3A shows sensitivity maps along with an indication of the sample position and shape. Every embryo was observed under a stereomicroscope before and after the NMR analysis to ensure positioning and verify successful sample retrieval with no physical damage.

**Figure 3:**
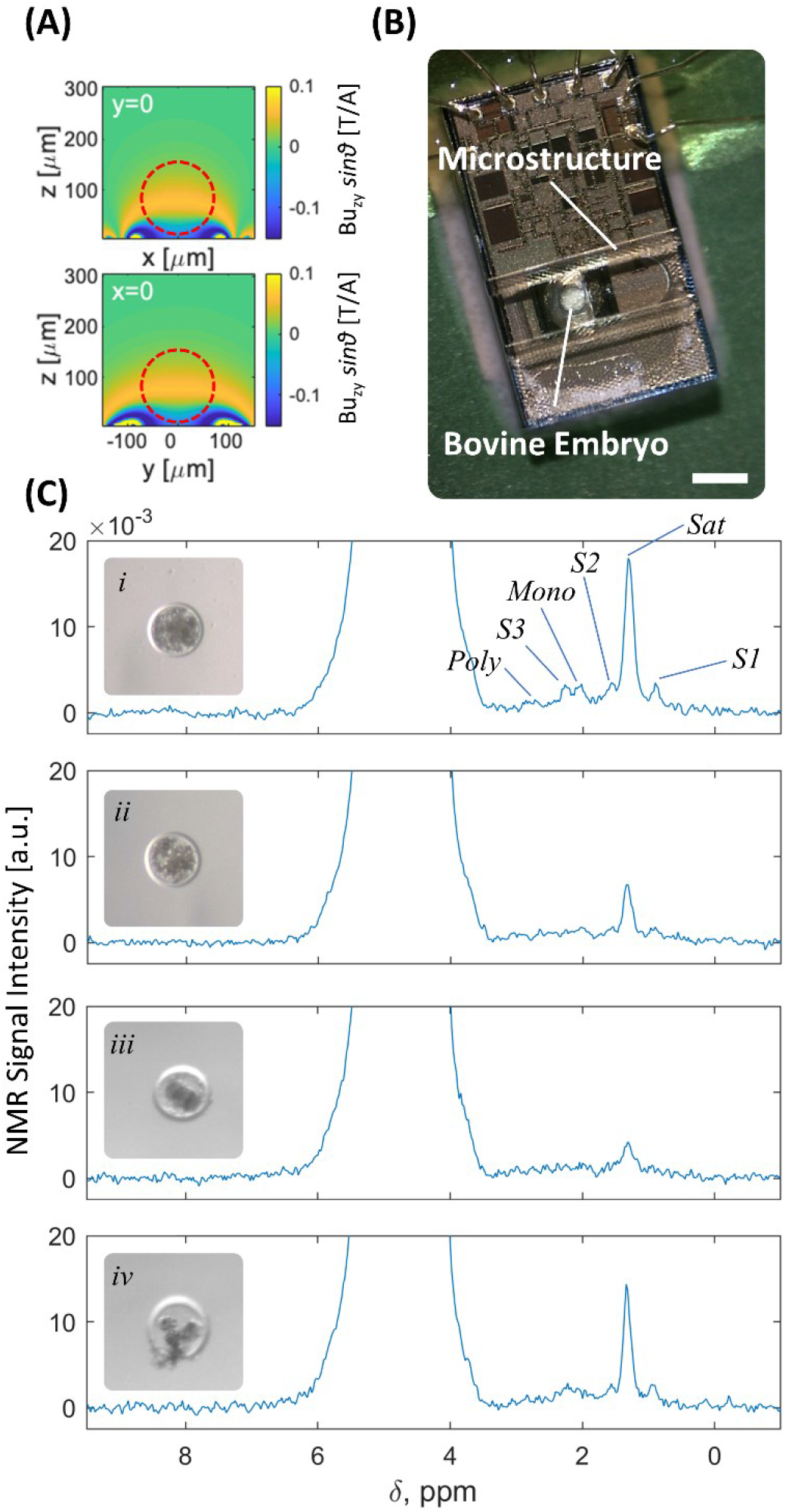
Micro-NMR spectroscopy of bovine embryos. **A)** Maps of sensitivity of the integrated microcoil in experimental conditions (*τ*=20 μs, *i*=11 mA) at *y*=0 and *x*=0 cross-sections. The static *B*_0_ field is oriented along the 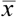 axis, while the 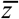 axis is perpendicular to the coil surface. The unitary field *Bu*_zy_, defined as the component orthogonal to *B*_*0*_ of the field produced by a current of 1 A in the coil loops, is computed via the Biot-Savart law. *Bu*_*zy*_ is also used to compute the nutation angle *θ*. Red dashed circular shapes indicate the sample position. **B)** Picture showing bovine embryo correct placement in the sample holding microstructure. Scale bar = 300 μm. **C)** Representative NMR spectra obtained from arrested early stage pre-implantation embryos and their corresponding photographs. Embryo diameters (μm) top to bottom: 187, 169, 137, 158. *(i)* and *(ii)*: arrested morulae, *(iii)* and *(iv)*: arrested 2-cell embryos.

Fig. 3C shows the NMR spectra of two morulae (*i* and *ii*) and two 2-cell stage arrested embryos (*iii* and *iv*). Despite the relatively large baseline-width of the water (∼ 2 ppm), the visible regions of the spectrum present a linewidth which is narrower to what previously observed in human microtissues (about 0.4 ppm)^31^ and similar to microorganism eggs^29,30^ (about 0.2 ppm). In these previous studies the field inhomogeneities are attributed to the presence of highly concentrated lipid droplets. In agreement with this hypothesis, these data would indicate that mammalian embryos have a lower concentration of lipid droplets with respect to human liver cell cultures. Similarly to previous studies, the linewidth cannot be improved with shimming when a sample is loaded, but better than 0.02 ppm FWHM are routinely obtained when the sensing region is occupied by a pure liquid medium^30^. This suggests that the origin of such relatively broad linewidths arises from magnetic susceptibilities mismatches within the sample. Randomly distributed lipid droplets, similarly to what is observed *in vivo*^37-39^, are a possible explanation the spectral resolution in our case.

The spectra present a significant heterogeneity among embryos, with up to 6 peak regions visible as indicated in Fig. 3C*i*. Table 1 compares their chemical shift to those of different molecular groups found in fatty acids in models of oils and organs extracts^37,40-42^. By comparison, the link to the sample fat content and in particular its fatty acids composition can be inferred (among which saturated, mono- and poly-unsaturated protons).

**Table 1:** Lipids chemical shifts. Chemical shift of lipids as reported in literature^37,40-42^ and comparison to bovine embryo spectra in Fig. 3C*i*.

| Chemical group | <sup>1</sup>H Shift ($\delta$ ppm) | Label (Fig. 3) |
| --- | --- | --- |
| CH<sub>3</sub>— | 0.83-1.03 | S1 |
| —(CH<sub>2</sub>)<sub>n</sub> | 1.22-1.42 | Sat |
| —CH<sub>2</sub>—CH<sub>2</sub>—CO<sub>2</sub>— | 1.52-1.70 | S2 |
| —CH=CH—CH<sub>2</sub>— | 1.94-2.14 | Mono |
| —CH<sub>2</sub>—CO<sub>2</sub>— | 2.23-2.36 | S3 |
| =CH—CH<sub>2</sub>—CH= | 2.70-2.84 | Poly |

The most visible heterogeneity is in the amplitude of the peaks, which is different within the same group (i.e., within the two morulae and within the two 2-cell stage arrested embryos). For example, the peak at 1.3 ppm has a variation of more than 200% in the two morulae samples. The same is true for the two early stage embryos. From the sensitivity map shown in Fig. 3A we estimated variations of the NMR signal due to sample volume variation. The expected signal change is 15% and 30% respectively comparing the two morulae and the two 2-cell embryos. Similar evaluations show that, with a precision in sample position of 20 μm, only 5% signal variation is expected. At last, thanks to the shape of the sensing region about half of the sample volume is observed through an ellipsoidal section crossing it approximately in the middle. In these conditions about half of the sample volume contributes to the signal and anisotropies in matter distribution are attenuated. The exaggerated approximation of the sample to a half-filled sphere shows only 25% expected signal variation upon a rotation of 180°. Furthermore, control experiments also indicated that the spurious effects resulting from sample positioning in the sensitive region of the micro coil do not explain the variations observed in their spectra.

As previously reported, liquid-state NMR spectroscopy will “bias” more mobile-lipids, i.e. those characterized by a certain degree of molecular tumbling^43^, as the tumbling improves line shape and thus detectability. This condition is more prominent in intracellular droplets having diameters larger than a few hundred nanometers allowing liquid-like tumbling in their interiors^44^. Lipid signals observed in comparative studies with Nile red staining and electron microscopy were confirmed to have origin in large lipid droplets, as further demonstrated in models of tumors and cell cultures^45-50^. They can be markers of apoptosis, necrosis, metabolic diseases, proliferation capabilities^43,48,51-53^.

## Discussion and Outlook

In a recent study on 3D human liver tissue cultures we demonstrated that spectra similar to those found here allowed for the definition of biomarkers for fatty liver disease^31^. We propose that the same method may be used also for mammalian embryos to monitor biologically fundamental processes. Generally, the majority of the analytical assays used in lipid metabolism investigations demand a large number of biological samples and rely on invasive/destructive methods such as histology and biopsies, which would hinder the continuation of embryonic development. A non-invasive *in vivo* assay would provide the means to reveal the role of embryonic lipids throughout development.

Arguably, the most important step toward future embryo applications and further establishment of NMR as a prime *in vivo* technique is the reliable detection of an expanded range of signals from cellular components. To further increase the information content several improvements can be implemented, especially on the NMR spectroscopy side, a few of which are summarized below.

### Improved Sensitivity and Water Suppression

as the sensitivity of NMR scales with a factor (ω/ω0)^7/454^ the simplest approach to improve sensitivity is to increase the magnetic field strength. For example, moving from 300 to 500 MHz (i.e. from 7 to 11 T) would increase the signal nearly 2.5 fold. While the use of the world’s largest commercial magnets (1.2 GHz) (28 T) would improve signal by a factor of 11. As SNR scales with √N the data that took 50 mins to collect here could probably be collected in less than 30 s at 1.2 GHz, potentially opening up the potential for close to real time molecular changes to be monitored *in vivo*. Another improvement would be the use of water suppression. The lack of pulsed field gradients on the setup used here negates cutting edge water suppression approaches^55,56^. Water suppression would reduce the water signal allowing the receiver gain to be increased.

### Multi-dimensional ^1^H NMR

Beyond sensitivity, the next hurdle is improving spectral dispersion from samples dominated by magnetic susceptibility broadening. The simplest way to achieve this would be via multidimensional homonuclear spectroscopy. Basic approaches such as ^1^H-^1^H COSY and TOCSY offer some potential *in vivo*, although spectral overlap and fast relaxation limit their resolving capabilities to only a handful of the most abundant compounds *in vivo*^9^. Conversely, dedicated approaches to reducing spectral inhomogeneities via intermolecular quantum interactions^57^ hold great potential. However, even the most sensitive of these approaches, termed In-phase Intermolecular Single Quantum Coherence (IP-ISQC)^58^, recovers only 10% of the net magnetization, which would require longer experimental times or higher magnetic fields for optimal analysis. Still, if successful, such approaches offer the potential to narrow and recover metabolites signals in an organism.

### Isotopic Enrichment and Heteronuclear nD NMR

While ^1^H NMR provides a theoretical peak capacity of around ∼3000 peaks, a 2D ^1^H-^13^C correlation increases this to ∼2 million, while 3D NMR approaches 100 million^59^. With such great improvements in spectral dispersion, even a moderate field of 500 MHz can be used to provide comprehensive assignment of a living metabolome^9^. However, such *in vivo* analysis generally requires isotopic enrichment (^13^C, or ^13^C and ^15^N) to increase signal due to the relative low biomass content of small biological entities. Such enrichment opens up many novel approaches including: amino-acid only NMR^60^ (essential as amino acids are involved in most stress and growth processes); moving window non-uniform sampling (close to real-time 2D NMR)^61^; and even isotope sub-editing which renders the organism “invisible” allowing only the ^13^C probe molecule (drug, toxin, nutrient) to be seen and monitored inside the organism^62^. However, while it is relatively easy to fully ^13^C enrich small aquatic species such *Hyalella azteca* (freshwater shrimp)^63^ and *Daphnia magna* (water fleas)^9^, it becomes near impossible to fully enrich cows or humans thus limiting this approach. This stated, Stable Isotope Resolved Metabolomics, where only specific pathways are targeted, holds great potential, and is making profound in-roads into understanding the complex processes behind complex diseases such as cancer^64^.

### Multiplexing

Beyond spectroscopic approaches, multiplexing (i.e. numerous CMOS NMR chips on the same device), could increase both throughput and sensitivity as-well as open up new applications^65^. This is possible as the homogeneous region of a modern narrow bore NMR magnet is relatively large (∼2 cm x 2 cm) and provides the space for a reasonable number of microchip-based NMR channels. At the most fundamental level multiple coils could allow the analysis of multiple eggs at the same time. However, it also opens up new possibilities especially those involving the monitoring of processes^15,66,67^, where control eggs and exposed eggs need to be monitored. Multiplexing different samples could allow to analyze control and exposed samples in parallel, reducing acquisition time and experimental variability. The complication of multiplexed approaches is the need to engineer complex micro-devices, eggs handling, and multiplexed fluidics required to keep arrays of eggs alive. To date, such development remains non-trivial.

## Conclusion

In this study we successfully utilized NMR spectroscopy (or MRS) on single mammalian embryos demonstrating the observation of fatty acids. The methodology shown can further develop into a non-invasive embryo assay for selection prior to embryo transfer. This would be applicable to both human and animal assisted reproductive techniques whose success rate today is fairly low, the current practice being largely based only on morphological scoring of the embryos. More in general, microchip NMR technology paves a new roadmap for the molecular analysis of microscopic biological entities. With applications in any area of research involving cells, eggs, or small organisms, the approach holds potential as a “next-generation” research tool across a diverse range of disciplines from environmental, through biochemical to medical.

## Author Contributions

GMC, MG, HB, CH conceived the study. CH, GS provided samples and optimized protocols. MG and GMC designed and fabricated the CMOS-based sensors. MG, GS, KM, and KJR performed experiments and data collection. GB, JB contributed to data interpretation and discussed micro-fabrication aspects. MG, KJR performed data analysis. MG supervised the project. MG, GS, AJS, GC wrote the manuscript. All authors edited and approved the content.

## Acknowledgements

We thank CPMA clinic (center of assisted reproduction, Lausanne) for providing culture incubators. We are grateful to the help of the staff at the Center of Micro and Nano Technology (CMi) at EPFL. This work was partially supported by Innosuisse under grant 39821.1 IP-LS and the European Union’s Horizon 2020 research and innovation program under grant agreement No 681002. AJS would like to thank the Natural Sciences and Engineering Research Council of Canada (Discovery (RGPIN-2019-04165), Alliance (ALLRP 549399) and Research Tools and Instruments (RTI-2020-00293)) Grant Programs

